# Evaluation of intranasal nafamostat or camostat for SARS-CoV-2 chemoprophylaxis in Syrian golden hamsters

**DOI:** 10.1101/2021.07.08.451654

**Authors:** M Neary, H Box, J Sharp, L Tatham, P Curley, J Herriott, E Kijak, U Arshad, JJ Hobson, RKR Rajoli, H Pertinez, A Valentijn, K Dhaliwal, F McCaughan, SP Rannard, A Kipar, JP Stewart, A Owen

## Abstract

Successful development of a chemoprophylaxis against SARS-CoV-2 could provide a tool for infection prevention implementable alongside vaccination programmes. Camostat and nafamostat are serine protease inhibitors that inhibit SARS-CoV-2 viral entry in vitro but have not been characterised for chemoprophylaxis in animal models. Clinically, nafamostat is limited to intravenous delivery and while camostat is orally available, both drugs have extremely short plasma half-lives. This study sought to determine whether intranasal dosing at 5 mg/kg twice daily was able to prevent airborne transmission of SARS-CoV-2 from infected to uninfected Syrian golden hamsters. SARS-CoV-2 viral RNA was above the limits of quantification in both saline- and camostat-treated hamsters 5 days after cohabitation with a SARS-CoV-2 inoculated hamster. However, intranasal nafamostat-treated hamsters remained RNA negative for the full 7 days of cohabitation. Changes in body weight over the course of the experiment were supportive of a lack of clinical symptomology in nafamostat-treated but not saline- or camostat-treated animals. These data are strongly supportive of the utility of intranasally delivered nafamostat for prevention of SARS-CoV-2 infection and further studies are underway to confirm absence of pulmonary infection and pathological changes.

## Introduction

Repurposing of previously approved drugs is an attractive strategy in the search for anti-SARS-CoV-2 treatment and prophylaxis. Certain sectors of society either cannot or will not benefit from the recent success with vaccines, and concerns around their longevity on the backdrop of new and future SARS-CoV-2 variants have been raised.^1^ Therefore, effective chemoprophylactic interventions represent a complimentary tool for deployment alongside national and international vaccination programmes. For other pathogens such as malaria, tuberculosis and HIV, successful prophylactic countermeasures have been developed using small molecule inhibitors of replication.^2-4^ The authors postulated that topical administration of an inhibitor of SARS-CoV-2 viral entry via intranasal delivery to healthy individuals may have utility in preventing transmission. Transmembrane protease serine 2 (TMPRSS2) is a protease found abundantly on the surface of cells within the respiratory tract^5^ and is utilised by the SARS-CoV-2 virus for S protein priming and activation, which enables virus entry into cells.^6^ TMPRSS2 activity is essential to viral pathogenesis of coronaviruses^7, 8^ and therefore presents putative opportunity as a drug target.

Camostat mesylate (camostat) and nafamostat mesylate (nafamostat) are serine protease inhibitors, used in the treatment of pancreatitis,^9, 10^ and have been demonstrated to bind and inhibit TMPRSS2.^11^ *In vitro* and *in vivo* studies of camostat, have demonstrated activity against SARS-CoV,^10-14^ with a study describing a 60% reduction in mortality following a lethal inoculum of SARS-CoV in mice receiving camostat at 30 mg/kg twice daily.^10^ More recently, topical camostat administered at 50 µM has been shown to block SARS-CoV-2 infection in human airway epithelial cells using an air-liquid interface model, indicating its potential use as prophylaxis against SARS-CoV-2.^15^ Furthermore, both camostat and nafamostat have been demonstrated to inhibit S-mediated entry of SARS-CoV-2 into lung cells *in vitro* with an approximately 15-fold higher potency of nafamostat compared to camostat.^14^ Nafamostat was also shown to block SARS-CoV-2 entry into Calu-3 cells with 10-fold higher potency than camostat.^11^

Nafamostat is also hypothesised to have a secondary effect upon thrombotic complications arising from COVID-19, which are markers of severe COVID-19 infection and linked to multi-organ failure and mortality.^16^ Nafamostat may inhibit platelet activation, resulting in subsequent inhibition of neutrophil extracellular traps and the contact factor activation pathway, which when activated alongside other pathways during SARS-CoV-2 infection may result in a prothrombotic state.^16^

The current study sought to assess the potential efficacy of intranasal nafamostat and camostat in preventing airborne acquisition of SARS-CoV-2 from infected to uninfected Syrian Golden Hamsters. The overarching aim was to provide preclinical data to support or refute utility of nafamostat or camostat as chemoprophylactic interventions for SARS-CoV-2.

## Methods

### Materials

Phosphate buffered saline (PBS) was purchased from Merck. Male Syrian Golden hamsters were purchased from Janvier Labs. 1 mL Amies Regular flocked swab were purchased from Appleton Woods. GoTaq^®^ Probe 1-Step RT-qPCR System was purchased from Promega. SARS-CoV-2 (2019-nCoV) CDC qPCR Probe Assay, CDC RUO 2019-nCoV_N_Positive Control and the SARS-CoV-2 E SgRNA were purchased from IDT. TRIzol reagent, GlycoBlue^™^, Phasemaker^™^ tubes, Nanodrop and TURBO DNA-free^™^ kit were purchased from ThermoFisher. A bead mill homogeniser was purchased from Fisher Scientific. Precellys CKmix lysing tubes were purchased from Bertin Instruments. A Chromo4^™^ Real-Time PCR Detector purchased from Bio-Rad. Transmission cages were purchased from Techniplast UK Ltd.

### Virus isolates

Human nCoV19 isolate/England/202012/01B (lineage B.1.1.7) was obtained from the National Infection Service at Public Health England, Porton Down, UK, via the European Virus Archive (catalogue code 004V-04032). This was supported by the European Virus Archive GLOBAL (EVA-GLOBAL) project that has received funding from the European Union’s Horizon 2020 research and innovation programme under grant agreement No 871029.

### Animal studies

All work involving SARS-CoV-2 was performed at containment level 3 by staff equipped with respirator airstream units with filtered air supply. Prior to the start of the study, all risk assessments and standard operating procedures were approved by the University of Liverpool Biohazards Sub-Committee and the UK Health and Safety Executive.

All animal studies were conducted in accordance with UK Home Office Animals Scientific Procedures Act (ASPA, 1986). Additionally, all studies were approved by the local University of Liverpool Animal Welfare and Ethical Review Body and performed under UK Home Office Project Licence PP4715265. Male Syrian golden hamsters (80-100 g; Janvier Labs) were housed in individually-ventilated cages with environmental enrichment under SPF barrier conditions and a 12-hour light/dark cycle at 21 °C ± 2 °C. Free access to food and water was provided at all times.

Hamsters were randomly assigned into groups of four and acclimatised for 7 days. Subsequently, three naïve hamsters in each group were intranasally dosed with 50 µL of saline (control), 5 mg/kg nafamostat in saline twice daily or 5 mg/kg camostat in saline twice daily. Following 24-hour cohabitation, the untreated hamster in each group was anaesthetised under 3% isoflurane and inoculated intranasally with 100 µL of 1 ⨯ 10^4^ PFU of nCoV19 isolate/England/202012/01B, lineage B.1.1.7 in phosphate buffered saline (PBS). These animals are henceforth referred to as the donor hamsters. Post-inoculation hamsters were housed in techniplast GR1800DIV cages with a divider that allows airflow from one side to the other (Figure 1). In each treatment group, the donor hamster was co-housed within the same cage as the naïve treated hamsters but physically separated by a plastic perforated barrier, to prevent contact transmission. The three naive hamsters in each group continued their respective dosing for a further 7 days post inoculation (day 0), before treatment was ended and directly inoculated hamsters were sacrificed. Treated hamsters were then housed for a further 7 days in the same cage until they were also sacrificed. All animals were weighed and monitored daily throughout the experiment, and throat swabs were taken on days 1, 3, 5, and 7. Naive hamsters were also swabbed at day 9, 11 and 14. In all cases, animal sacrifice was conducted via a lethal intraperitoneal injection of pentobarbitone, followed by cardiac puncture and immediate exsanguination of blood from the heart.

**Figure 1:**
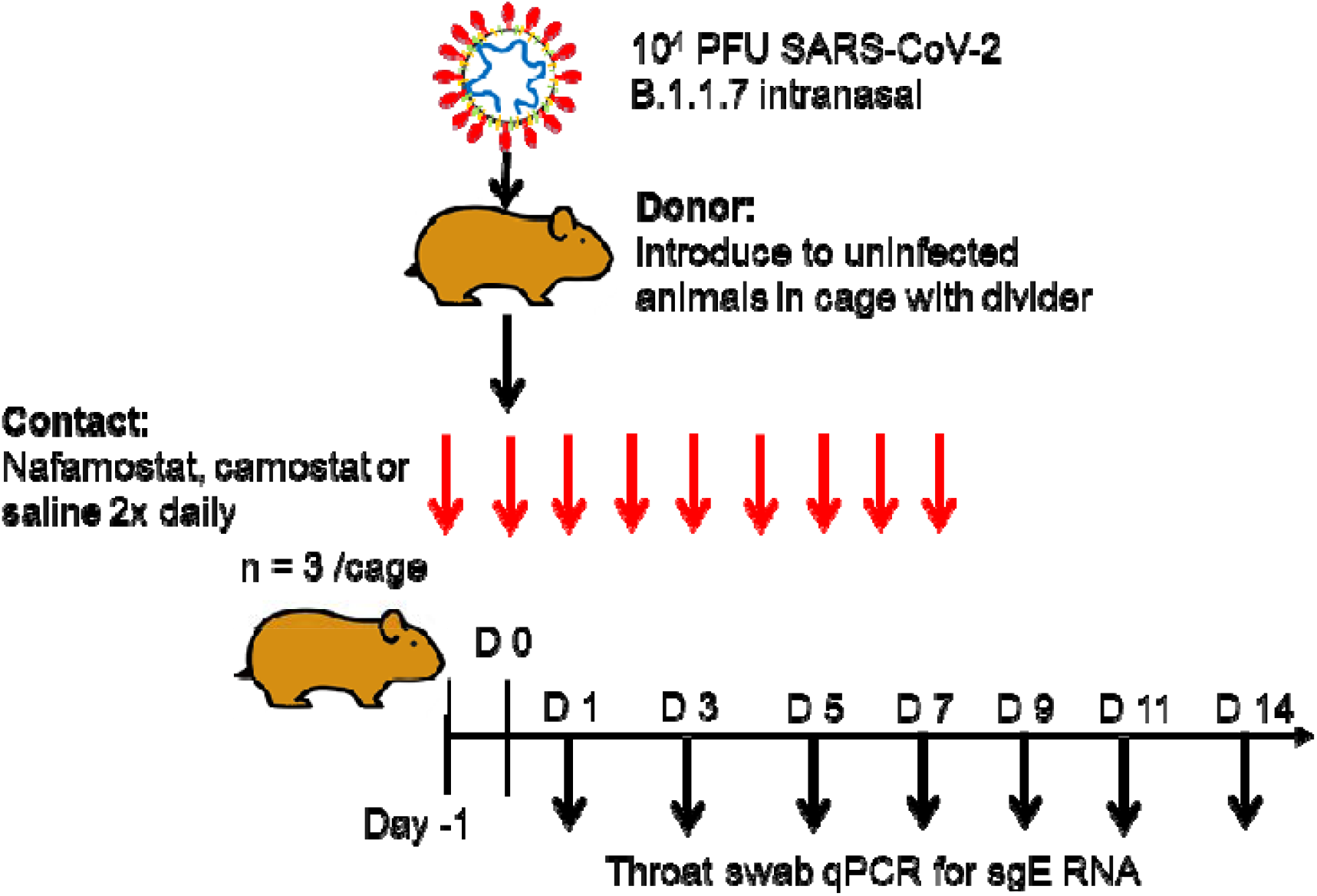
Diagrammatic representation of the experimental design employed to assess chemoprophylaxis.

### Quantification of viral RNA

A section of dissected lung lobe and nasal turbinate material was homogenised in 1 mL of TRIzol reagent (ThermoFisher) using a bead mill homogeniser (Fisher Scientific) and Precellys CKmix lysing tubes (Bertin Instruments) at 3.5 metres per second for 30 seconds. The resulting lysate was centrifuged at 12,000 x g for 5 min at 4^°^C. Throat swab media (260 µL) was added to 750 µL of TRIzol LS reagent (ThermoFisher). The clear supernatants were transferred to Phasemaker^™^ tubes (ThermoFisher) and processed as per the manufacturer’s instructions to separate total RNA from the phenol-chloroform layer. Subsequently, the recovered RNA was precipitated using GlycoBlue^™^ according to the manufacturer’s instructions (ThermoFisher), washed and solubilised in RNAse-free water. The RNA was quantified and quality assessed using a Nanodrop (ThermoFisher). Samples were diluted to either 20,000 or 200 ng/mL in 60 µL of RNAse-free water. The resulting RNA samples were DNAse treated using the TURBO DNA-free^™^ kit according to the manufacturer’s instructions (ThermoFisher). The DNAse treated RNA was stored at -80^°^C prior to downstream analysis.

The viral RNA derived from hamster lung, nasal turbinate and throat swabs was quantified using a protocol adapted from the CDC 2019-Novel Coronavirus (2019-nCoV) Real-Time PCR Diagnostic Panel^17^ and a protocol for quantifying the SARS-CoV-2 subgenomic E gene RNA (E SgRNA)^18^ using the GoTaq^®^ Probe 1-Step RT-qPCR System (Promega). For quantification of SARS-CoV-2 using the nCoV assay, the N1 primer/probe mix from the SARS-CoV-2 (2019-nCoV) CDC qPCR Probe Assay (IDT) were selected. A standard curve was prepared (1,000,000 – 10 copies/reaction) via a 10-fold serial dilution of the CDC RUO 2019-nCoV_N_Positive Control (IDT). DNAse treated RNA at 200 ng/mL or dH_2_O was added to appropriate wells producing final reaction volumes of 20 µL. The prepared plates were run using a Chromo4^™^ Real-Time PCR Detector (Bio-Rad). The thermal cycling conditions for the qRT-PCR reactions were: 1 cycle of 45^°^C for 15 min, 1 cycle of 95^°^C for 2 min, f°llowed by 45 cycles of 95^°^C for 3 seconds and 55^°^C for 30 seconds.

Quantification of SARS-CoV-2 E SgRNA was completed utilising primers and probes previously described elsewhere^18^ and were used at 400 nM and 200 nM, respectively (IDT), using the GoTaq^®^ Probe 1-Step RT-qPCR System (Promega). Quantification of 18S RNA utilised previously described primers and probe sequences^19^, and were used at 300 nM and 200 nM, respectively (IDT), using the GoTaq^®^ Probe 1-Step RT-qPCR System (Promega). Methods for the generation of the 18S and E SgRNA standards have been outlined previously.^20^ Both PCR products were serially diluted to produce standard curves in the range of 5 × 10^8^ - 5 copies/reaction via a 10-fold serial dilution. DNAse treated RNA at 20,000 ng/mL or dH_2_O were added to appropriate wells producing final reaction volumes of 20 µL. The prepared plates were run using a Chromo4^™^ Real-Time PCR Detector (Bio-Rad). The thermal cycling conditions for the qRT-PCR reactions were: 1 cycle of 45^°^C for 15 min, 1 cycle of 95^°^C for 2 min, followed by 45 cycles of 95^°^C for 3 seconds and 60^°^C for 30 seconds. Both N and E SgRNA data were normalised to 18S data for subsequent quantitation.

### Statistical analysis

An unpaired t-test was used to compare the differences in body weight compared to that observed at baseline in the saline control group at day 8, and the nafamostat or camostat treatment groups at day 8. A *P* value of ≤ 0.05 was taken as statistically significant. All statistical analysis was completed using GraphPad Prism version 8.3.0.

## Results

### Infection-mediated changes in body weight

To test the efficacy of respiratory administration of nafamostat or camostat against SARS-CoV-2 infection, hamsters were treated for 24 h before being exposed to infected hamsters in cages with perforated dividers to mimic non-contact, respiratory transmission as detailed in Figure 1. Contact hamsters were treated daily for a further 7 days and then maintained for a further 7 days without treatment.

Body weight changes in each group over the course of the experiment are shown in Figure 2. Saline-treated hamsters in the control group increased in weight by an average of 5% from day -1 to day 4 after infection of inoculated animals, before declining in weight by 11% from day 4 through to day 9. Body weight then increased by 11.5% between day 9 and day 13.

**Figure 2:**
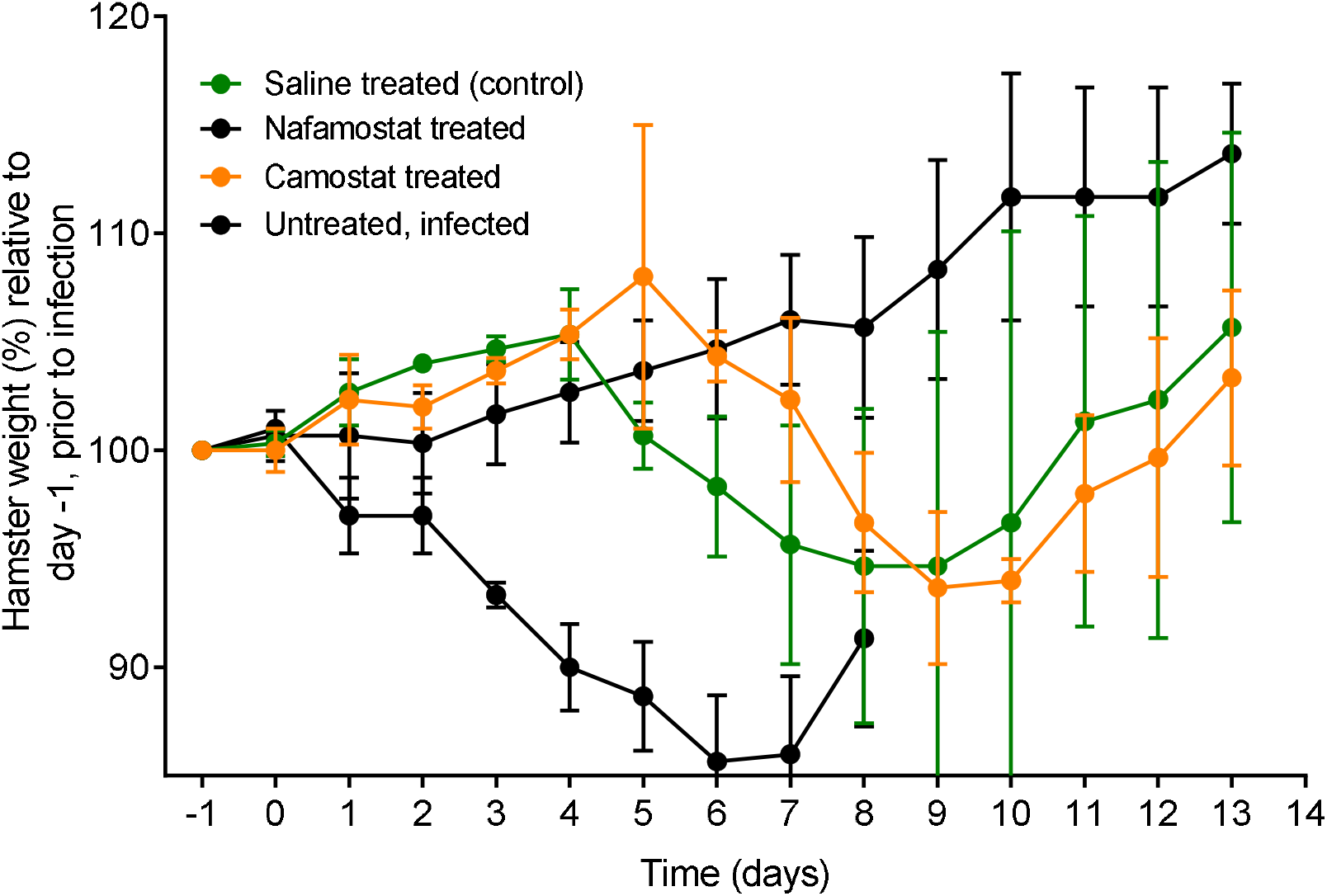
Hamster weights separated by group and inoculation status over the study time course. All hamsters within each treatment group (n = 3), and the donor hamster cohoused with each group, were weighed at 24 hourly intervals up to their study endpoint. All weights are shown as a percentage of the initial weight recorded at baseline, day -1 of the study.

Nafamostat-treated hamsters increased in weight by an average of 13.3% compared to their original day -1 mean weight, with no decrease in mean weight observed throughout the study. The difference in mean weight change compared to that observed at baseline on day -1 in the nafamostat-treated hamsters, compared to the saline treated control group at day 8, was seen to be statistically significant (*P* = 0.004).

Within the camostat group, average weight was seen to increase up to day 5 before declining by 5% over a 6-day period to 99.2% of the day -1 weight. The average weight at day 13 was 3.4% higher than the baseline weight recorded on day -1. The average weight change at day 8 in the camostat group, was not seen to be statistically significant to that observed within the saline treated (control) group (*P* = 0.105).

### Viral RNA measurements in swab samples during co-habitation

The quantification of virus subgenomic RNA specific for the E gene (sgE) from throat swabs are shown in Figure 3. Subgenomic RNAs are only produced during virus replication and so are an indication of active infection. For saline-treated controls, the mean SARS-CoV-2 sgE RNA concentration from the swabs rose above the LOQ for the assay to 1,100 copies of sgE RNA/µg of RNA relative to 18S, at day 5. This then declined to 41.5 copies of sgE RNA/µg of RNA relative to 18S at day 7. Indicating an establishment of infection in the saline treated group from day 5 of the study whilst cohoused with the donor hamster.

**Figure 3:**
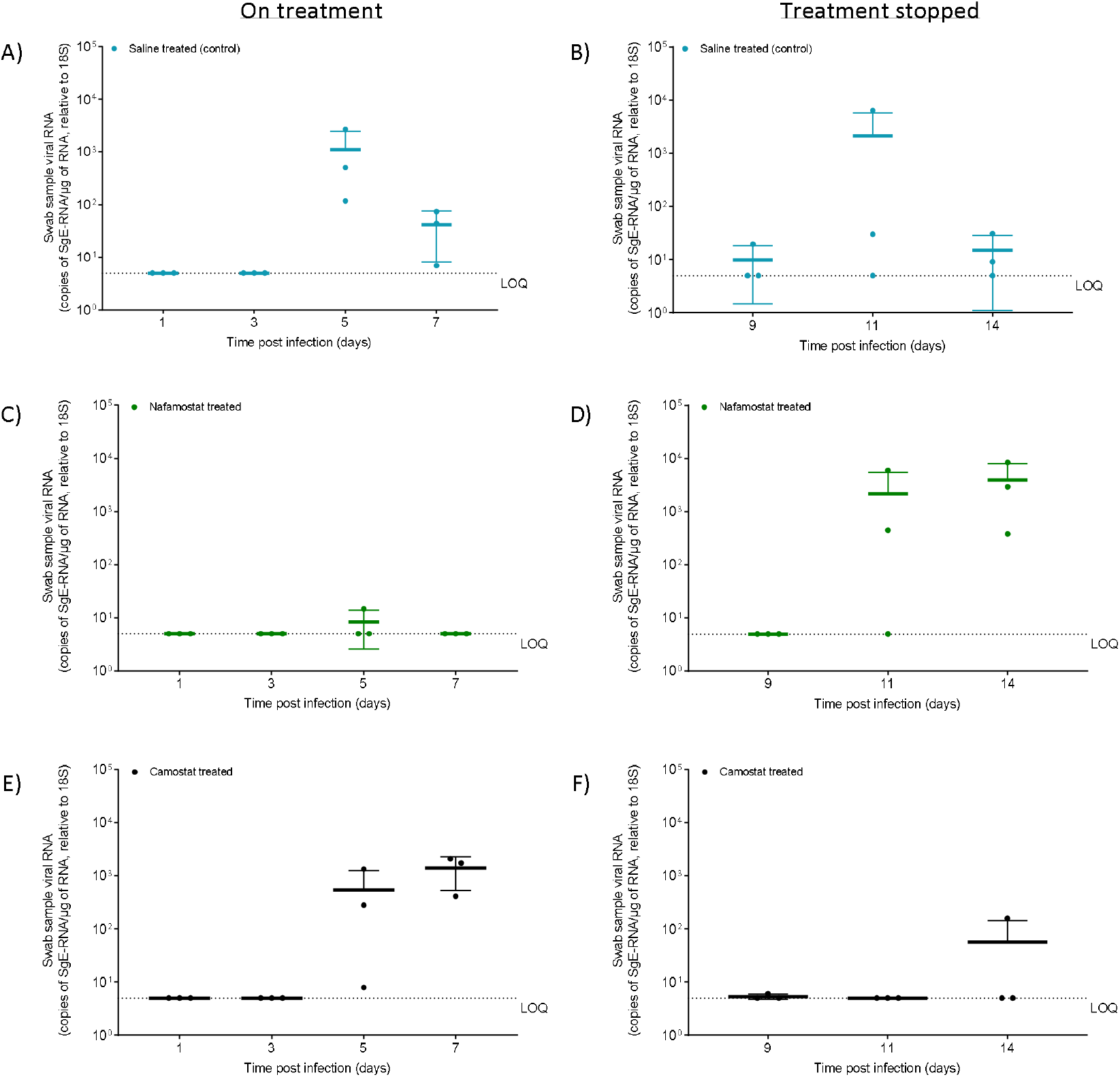
Viral quantification of SARS-CoV-2 sgE RNA within swab samples obtained from the treated groups up to day 14 of the study. Viral quantification of SARS-CoV-2 sgE RNA within swab samples from A) the saline treated (control) hamsters (n = 3) during treatment, B) the saline treated (control) hamsters (n = 3) after stopping treatment, C) the nafamostat-treated hamsters (n = 3) during treatment, D) the nafamostat-treated hamsters (n = 3) after stopping treatment, E) the camostat-treated hamsters (n = 3) during treatment, F) the camostat-treated hamsters (n = 3) after stopping treatment.

For the nafamostat-treated group the average SARS-CoV-2 sgE RNA from the swabs was observed to be below the LOQ up to day 7, with the exception of day 5 at 5.0 copies of sgE RNA/µg of RNA, relative to 18S driven by low levels in one animal. This demonstrated a lack of detectable virus in the nafamostat treated group during the 7 days of nafamostat treatment and cohousing with the donor hamster.

The swab samples taken from the camostat treated group had an average SARS-CoV-2 sgE RNA concentration above the LOQ from day 5 onwards. A peak viral load of 2,090 copies of sgE RNA/µg of RNA relative to 18S was reached at day 7. This data indicates the establishment of SARS-CoV-2 infection in all of the camostat treated hamsters after day 5, during the camostat treatment and cohousing with the donor hamster.

At day 9 the saline-treated controls had an average SARS-CoV-2 sgE RNA swab concentration of 8.1 copies of sgE RNA/µg of RNA, relative to 18S. Increasing at day 11 to 2,126 copies of sgE RNA/µg of RNA relative to 18S, before subsequently declining to 14.8 copies of sgE RNA/µg of RNA relative to 18S, at day 14. Indicating viral clearance in the saline treated control group after termination of cohousing with the donor hamster.

Within the nafamostat-treated group virus concentrations continued to be below the LOQ at day 9. However, at day 11 the average concentration sgE RNA/µg of RNA relative to 18S was observed to be 2,138 copies, followed by 3,907 copies at day 14. This indicates establishment of a detectable viral infection in the nafamostat treated hamsters 4 days post termination of cohousing with the donor hamster and cessation of nafamostat treatment.

From day 9 onwards the camostat-treated animals had an average SARS-CoV-2 sgE RNA swab concentration below the LOQ up to day 14, where a single hamster had a detectable viral load of 159 copies of sgE RNA/µg of RNA, relative to 18S. Indicating viral clearance from day 9 onwards, 48 hours after cohousing with the donor housing and camostat treatment ended.

## Discussion

This study sought to determine the suitability of intranasally delivered nafamostat or camostat for use as chemoprophylaxis for SARS-CoV-2 in an airborne transmission model in Syrian golden hamsters. Intranasal infection of Golden Syrian hamsters with SARS-CoV-2 has previously been demonstrated to result in viral titres and pathology similar to that seen within humans.^21^ Furthermore, *In silico* modelling has demonstrated the immune activation and cardiovascular changes in hamsters observed within 14 days post infection (dpi) mimicked the immediate COVID-19 pathologies observed within humans.^22^ Changes in the metabolomics profile within the hamsters was also seen to correlate with the alterations observed within the severe patient’s metabolomics profile.^22^ Importantly for the current study, TMPRSS2-mediated priming of the SARS-CoV-2 S protein was found to be largely similar in hamsters and humans,^22^ further highlighting the similarity between the host-pathogen response in hamsters and humans and supporting the use of this model for TMPRSS2-targeted therapeutics. However, it should be noted that hamsters are obligate nasal breathers,^23^ which presents a potential limitation of this species for characterising intranasally delivered chemoprophylactic interventions. The importance of this for future applicability in humans is uncertain but it should be noted.

Detectable viral RNA in swab samples from day 1 through day 7 were observed in animals receiving intranasal camostat, indicating a lack of chemoprophylactic benefit against SARS-CoV-2 infection at a dose of 10mg/kg/day. Conversely, viral RNA in swabs collected from the intranasal nafamostat group remained below the LOQ throughout the period of cohabitation with infected animals. Importantly, body weight has emerged as an important clinical outcome for severity of disease in the hamster model.^24^ Nafamostat but not camostat was also able to prevent the infection-mediated body weight decline that was observed in both directly inoculated and saline-treated control animals. Taken collectively, these data clearly demonstrate that 5mg/kg of intranasally twice daily-administered nafamostat but not camostat is able to prevent transmission from infected animals to healthy animals while they receive the drug.

Viral RNA became detectable within the throat swabs of nafamostat-treated animals on day 9, two days after discontinuation of the drug and despite infected animals being culled at the time of drug discontinuation. The current experimental design is not suitable to determine the reasons why these animals became infected after the directly inoculated animals had been sacrificed but it is possible that infection occurred via residual infectious virus present within the cage or within the nasal cavity of the animals. These data do however suggest that protection by nafamostat is short lived following discontinuation of intranasal dosing, with RNA becoming detectable 2 days following discontinuation, which is similar in duration to the initial detectability in saline-treated controls. In camostat-treated animals, the viral RNA was below the LOQ in most animals from day 9 onwards, presumably because the initial infection that occurred during dosing had then been cleared. A single hamster did however have detectable viral RNA at day 14 within the camostat group.

Intranasal dosing of test therapeutics enables direct drug delivery to a primary site of SARS-CoV-2 infection which may be particularly useful for chemoprophylaxis. However, it is currently unclear whether nafamostat delivered via this route would be sufficient to meaningfully alter the course of disease if given in a therapy model. A study of 420 µg/mL of nafamostat suspended within a novel lipid formulation and administered intranasally to hamsters, who were immediately inoculated with SARS-CoV-2, showed a transient but significant reduction in viral load within the nasal cavity compared to control. However, no impact on pathology was reported.^25^ It is not currently possible to ascertain whether the limited benefit were a result of the formulation itself since unformulated nafamostat was not used as a control in the published study. However, studies in a primary lung epithelium cell model did demonstrate low cytotoxicity of the lipid formulation up to 6 µg/mL nafamostat and evidence of SARS-CoV-2 inhibition at this dose.^26^

Given that the systemic half-life of nafamostat after intravenous delivery is only 8 minutes,^27^ the protection reported herein with twice daily administration bodes extremely well for the residence of nafamostat at its target when intranasally administered. Nafamostat is not currently approved for oral or inhalational administration in humans so development of a specific formulation may be needed if this approach is to be tested in humans. Further work by the authors is ongoing, to ensure reproducibility of these observations and to confirm an absence of pulmonary virus and pathology in hamsters given nafamostat as chemoprophylaxis. This preprint will be updated when these data become available.

In conclusion, a protective effect of 5 mg/kg twice daily intranasal dosing of nafamostat is reported that was not seen for camostat at the same dose. If these findings prove to be reproducible in subsequent studies, clinical evaluation of intranasal nafamostat safety may be warranted to enable larger trials as a chemoprophylaxis for COVID-19.

## Acknowledgements

Human nCoV19 isolate/England/202012/01B (lineage B.1.1.7) was provided by the National Infection Service at Public Health England, Porton Down UK via the European Virus Archive (catalogue code 004V-04032). This was supported by the European Virus Archive GLOBAL (EVA-GLOBAL) project that has received funding from the European Union’s Horizon 2020 research and innovation programme under grant agreement No 871029.

## References

1. Kuzmina A, Khalaila Y, Voloshin O et al. SARS-CoV-2 spike variants exhibit differential infectivity and neutralization resistance to convalescent or post-vaccination sera. Cell host & microbe 2021; 29: 522–8. e2.

2. Li H, Van Der Linden WA, Verdoes M et al. Assessing subunit dependency of the Plasmodium proteasome using small molecule inhibitors and active site probes. ACS chemical biology 2014; 9: 1869–76.

3. Godbole AA, Ahmed W, Bhat RS et al. Targeting Mycobacterium tuberculosis topoisomerase I by small-molecule inhibitors. Antimicrobial agents and chemotherapy 2015; 59: 1549–57.

4. Christ F, Shaw S, Demeulemeester J et al. Small-molecule inhibitors of the LEDGF/p75 binding site of integrase block HIV replication and modulate integrase multimerization. Antimicrobial agents and chemotherapy 2012; 56: 4365–74.

5. Bittmann S, Luchter E, Weissenstein A et al. TMPRSS2-inhibitors play a role in cell entry mechanism of COVID-19: an insight into camostat and nafamostat. J Regen Biol Med 2020; 2: 1–3.

6. Hoffmann M, Kleine-Weber H, Schroeder S et al. SARS-CoV-2 cell entry depends on ACE2 and TMPRSS2 and is blocked by a clinically proven protease inhibitor. cell 2020; 181: 271-80. e8.

7. Iwata-Yoshikawa N, Okamura T, Shimizu Y et al. TMPRSS2 contributes to virus spread and immunopathology in the airways of murine models after coronavirus infection. Journal of virology 2019; 93.

8. Shirato K, Kawase M, Matsuyama S. Wild-type human coronaviruses prefer cell-surface TMPRSS2 to endosomal cathepsins for cell entry. Virology 2018; 517: 9–15.

9. Takahashi W, Yoneda T, Koba H et al. Potential mechanisms of nafamostat therapy for severe COVID-19 pneumonia with disseminated intravascular coagulation. International Journal of Infectious Diseases 2021; 102: 529–31.

10. Zhou Y, Vedantham P, Lu K et al. Protease inhibitors targeting coronavirus and filovirus entry. Antiviral research 2015; 116: 76–84.

11. Yamamoto M, Kiso M, Sakai-Tagawa Y et al. The anticoagulant nafamostat potently inhibits SARS-CoV-2 S protein-mediated fusion in a cell fusion assay system and viral infection in vitro in a cell-type-dependent manner. Viruses 2020; 12: 629.

12. Hoffmann M, Hofmann-Winkler H, Smith JC et al. Camostat mesylate inhibits SARS-CoV-2 activation by TMPRSS2-related proteases and its metabolite GBPA exerts antiviral activity. EBioMedicine 2021; 65: 103255.

13. Kawase M, Shirato K, van der Hoek L et al. Simultaneous treatment of human bronchial epithelial cells with serine and cysteine protease inhibitors prevents severe acute respiratory syndrome coronavirus entry. Journal of virology 2012; 86: 6537–45.

14. Hoffmann M, Schroeder S, Kleine-Weber H et al. Nafamostat mesylate blocks activation of SARS-CoV-2: new treatment option for COVID-19. Antimicrobial agents and chemotherapy 2020; 64.

15. Guo W, Porter LM, Crozier TW et al. Topical TMPRSS2 inhibition prevents SARS-CoV-2 infection in differentiated primary human airway cells. bioRxiv 2021.

16. McFadyen JD, Stevens H, Peter K. The emerging threat of (micro) thrombosis in COVID-19 and its therapeutic implications. Circulation research 2020; 127: 571–87.

17. Wen C, Yee S, Liang X et al. Genome-wide association study identifies ABCG2 (BCRP) as an allopurinol transporter and a determinant of drug response. Clinical Pharmacology & Therapeutics 2015; 97: 518–25.

18. Wölfel R, Corman VM, Guggemos W et al. Virological assessment of hospitalized patients with COVID-2019. Nature 2020; 581: 465–9.

19. Ashraf N, Zino S, Macintyre A et al. Altered sirtuin expression is associated with node-positive breast cancer. British journal of cancer 2006; 95: 1056–61.

20. Salzer R, Clark JJ, Vaysburd M et al. Single-dose immunisation with a multimerised SARS-CoV-2 receptor binding domain (RBD) induces an enhanced and protective response in mice. Cold Spring Harbor Laboratory, 2021.

21. Imai M, Iwatsuki-Horimoto K, Hatta M et al. Syrian hamsters as a small animal model for SARS-CoV-2 infection and countermeasure development. Proceedings of the National Academy of Sciences 2020; 117: 16587–95.

22. Rizvi ZA, Dalal R, Sadhu S et al. Immunological and cardio-vascular pathologies associated with SARS-CoV-2 infection in golden syrian hamster. bioRxiv 2021.

23. Kling MA. A review of respiratory system anatomy, physiology, and disease in the mouse, rat, hamster, and gerbil. The veterinary clinics of North America Exotic animal practice 2011; 14: 287.

24. Rosenke K, Meade-White K, Letko M et al. Defining the Syrian hamster as a highly susceptible preclinical model for SARS-CoV-2 infection. Emerging Microbes & Infections 2020; 9: 2673-84.

25. Cornelissen L, Hoefsmit E, Rao D et al. Nafamostat Mesylate in lipid carrier for nasal SARS-CoV2 titer reduction in a hamster model. bioRxiv 2020.

26. Kirkpatrick L, Millard J. Evaluation of nafamostat mesylate safety and inhibition of SARS-CoV-2 replication using a 3-dimensional human airway epithelia model. bioRxiv 2020.

27. Okajima K, Uchiba M, Murakami K. Nafamostat mesilate. Cardiovascular Drug Reviews 1995; 13: 51–65.

